# Characterization of non-monotonic relationships between tumor mutational burden and clinical outcomes

**DOI:** 10.1101/2024.01.16.575937

**Authors:** Jordan Anaya, Julia Kung, Alexander S. Baras

## Abstract

Potential clinical biomarkers are often assessed with Cox regressions or their ability to differentiate two groups of patients based on a single cutoff. However, both of these approaches assume a monotonic relationship between the potential biomarker and survival. Tumor mutational burden (TMB) is currently being studied as a predictive biomarker for immunotherapy, and a single cutoff is often used to divide patients. In this study we introduce a two-cutoff approach that allows splitting of patients when a non-monotonic relationship is present, and explore the use of neural networks to model more complex relationships of TMB to outcome data. Using real-world data we find that while in most cases the true relationship between TMB and survival appears monotonic, that is not always the case and researchers should be made aware of this possibility.

**Significance:** When a non-monotonic relationship to survival is present it is not possible to divide patients by a single value of a predictor. Neural networks allow for complex transformations and can be used to correctly split patients when a non-monotonic relationship is present.

## INTRODUCTION

When searching for features predictive of survival across different cancer types researchers often use Cox regression^1–3^. The Cox model provides a high level of interpretability in the form of the regression coefficients, but these coefficients simply describe the linear relationship of the feature to the predicted log partial hazard^4^. Often there isn’t a clear reason to assume a linear (and more importantly monotonic) relationship; furthermore, neural nets have previously been proposed as a more flexible approach in this context^5^, with some recent applications appearing in the literature^6–9^.

Tumor mutational burden (TMB) is a commonly investigated biomarker in the context of immunotherapy^10–16^, and its prognostic value has also been investigated in the context of heterogeneous treatments^17,18^. When investigating TMB as a biomarker researchers often bin patients into a “TMB low” group and a “TMB high” group, which implicitly assumes a monotonic relationship of TMB with survival, independent of the number of bins used. In this case the relationship is assumed to be a step function with patients below a certain threshold having a certain risk and patients above the threshold having another. The monotonic assumption is that change in risk only increases (or only decreases) with the value of the predictor variable in question. However, other types of relationships could easily be envisioned. For example, while an increase in mutations may initially increase the pathogenicity of a tumor, too many mutations may be detrimental for tumor growth. Alternatively, high or low values of TMB could be a proxy for a different population of samples, such as high TMB being associated with microsatellite instability.

Given enough parameters and data, neural networks are able to learn arbitrarily complex functions. Similar to how we previously leveraged this flexibility to more optimally model the calibration of next-generation sequencing (NGS) gene panel-derived estimates of exomic TMB^19^, we wondered whether such flexible modeling approaches could be applied to characterizing the relationship of TMB with clinical outcomes data. While a single-cutoff approach may work well for monotonic relationships, it would be expected to be suboptimal in the case of a truly non-monotonic relationship. In such a scenario the lowest and highest TMB values would have similar risk, and the moderate values would be associated with a different risk. In this study we explored different approaches to attempt to better characterize these more complex relationships and investigated whether such relationships exist in the context of TMB and clinical outcomes data, both in the prognostic sense and predictive sense.

## Materials and Methods

Code for reproducing the results in this manuscript is available at https://github.com/OmnesRes/tmb survival and has been archived at Zenodo: https://doi.org/10.5281/zenodo.10966684.

### Modeling

Cox-PH models were implemented with lifelines version 0.27.7^20^ and the neural nets with TensorFlow 2.12^21^. TMB values were log transformed before inclusion into the models.

Looking at the formula for hazard in the Cox proptional hazards model it is easy to see that it is a function of a linear combination of the predictors^4^:

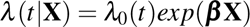

The hazard for a given set of predictors **X** (*X*_0_, *X*_1_, …) at time *t* is given by a baseline hazard *λ*_0_(*t*) and a partial hazard *exp*(***β*****X**). The log partial hazard is then ***β*X** (*β*_0_*X*_0_ + *β*_1_*X*_1_…). In our case the only predictor is TMB so the log partial hazard is given by *β*_*TMB*_*TMB*. In order for the partial hazard to be a more complex function of TMB the variable would have to undergo a transformation prior to inclusion into a Cox model, but the transformation that should be performed is unknown. For an example of a simple polynomial transformation see Supplemental Figure 1.

For the neural nets we interpret the output to be the log partial hazard, and use it to calculate the negative partial likelihood, as previously proposed^5^. The neural net is essentially a universal function approximator, and it transforms the input variable (TMB) into a risk score for each sample. The neural nets were designed to have two dense layers of 128 with softplus activation and a dropout of .05, followed by a dense to 1 with no activation. A batch normalization layer was utilized to keep the output values centered at 0.

When training with the real-world datasets the TMB values in the top 1% were discarded to avoid training with extreme values. Stratified K-fold training was performed with 10 train/test splits and stratified by whether the TMB value was in the top 20th percentile to ensure a somewhat consistent range of TMB values across training splits. For each fold the ranks of the test fold data were recorded in order to calculate the concordance indexes of the models. To ensure a comparable loss calculation between the neural net and Cox models lifelines was used to calculate all losses, with the predicted risks from the neural nets being passed into a Cox model from lifelines.

For the cutoff analyses we found the optimal cutoffs as determined by logrank statistic by exhaustively searching every possible data split. For the single-cutoff analyses we required each group to have at least 25 samples in it, and for the double cutoff analyses we required the moderate group to have at least 50 samples and the low and high groups to each have at least 25 samples.

### Data Processing

Simulated survival data was generated by utilizing an exponential distribution^22^, and a uniform distribution was used for censoring with approximately 30% of the data censored. For the simulated data only TMB values below 64 were used since it was difficult to prevent extreme simulated risks for our quadratic relationship, which highlights a potential issue of using polynomials for modeling data with a long tail.

When generating the log partial hazards for the simulated data TMB values were first log transformed. For the linear simulated data the risks (log partial hazards) were set to .5 *× TMB*. For the non-monotonic data we set the risks to (*exp*(*−*(2*TMB−* 1.5)^2^*/*(4*TMB*)) *−*.6) *×* 2. For the quadratic data we set the risks to ((*−*(*TMB−* 2)^2^) *×* 15 + 40) *×*.05. For the step data if the TMB value was below the median TMB value then the risk was set to 0, otherwise it was set to 1.

The TCGA somatic mutation calls were processed as previously described to calculate exomic TMB^19^. We used all available panels in the BPC data except for VICC-01-SOLIDTUMOR and UHN-48-V1 due to their small sizes. We used GENIE release 14.1^23^ to obtain somatic mutations and panel coordinates, and defined TMB as nonsynonymous mutations per Mb of panel coding sequence. PyRanges^24^ was used to perform intersections. For BPC we only included patients whose pathology procedure occurred within 180 days of diagnosis.

### Data Availability Statement

Publicly available data generated by others were used by the authors. TCGA somatic calls^25^ were obtained from https://gdc.cancer.gov/about-data/publications/mc3-2017. TCGA clinical data was obtained from Liu et al^26^. MSK datasets were obtained from Samstein et al. and Valero et al^15,18^. BPC data is available at https://www.synapse.org/#!Synapse: syn27056700. GENIE 14.1 panel information was obtained from https://www.synapse.org/#!Synapse:syn52918985. The GFF3 is available at ftp://ftp.ensembl.org/pub/grch37/current/gff3/homosapiens/Homosapiens.GRCh37.87.gff3.gz. The Broad coverage WIGs are available at https://www.synapse.org/#!Synapse:syn21785741.

## RESULTS

Given the long right-tailed distribution of TMB (Figure 1A), when modeling the relationship of this data to the log partial hazard some form of a transformation will generally be required, since these extreme values multiplied by the estimated model coefficient will generate unrealistic partial hazards. We can demonstrate this by fitting a Cox model to untransformed TMB values for uterine corpus endometrial carcinoma (UCEC) and the corresponding survival data in the The Cancer Genome Atlas (TCGA) dataset^26^. Generating predicted survival curves for several different TMB values (Figure 1B), we see that what the field would consider large differences in TMB result in minimal differences in the curves, and at the highest TMB values patients are predicted to survive longer than humanly possible.

**Figure 1.**
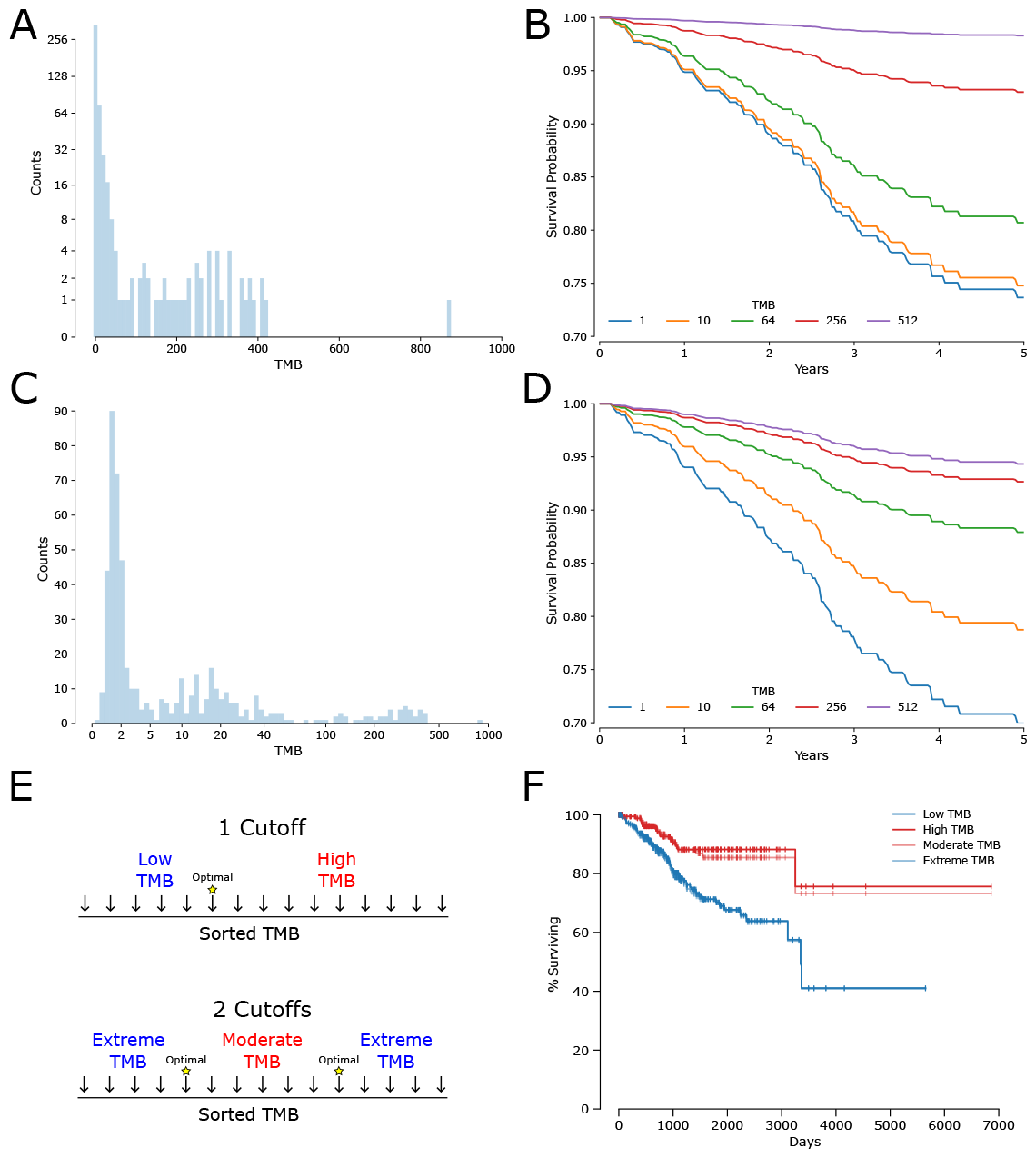
TMB transformation strategies. A, the TMB distribution for the TCGA UCEC data. B, predicted survival curves for several TMB values for a univariate Cox model. C, distribution for a log transformation of the TMB values. D, the resulting survival curves for a univariate Cox model with the log-transformed values. E, two strategies for creating a binary label based on TMB, a single-cutoff approach and a two-cutoff approach. F, the resulting survival curves for the most optimal cutoffs with both binary label approaches.

One solution to an extreme variable distribution is to log transform the data (Figure 1C). We see that fitting a Cox model to this transformed data now generates distinct survival curves for TMB values 1 and 10 (Figure 1D), but the survival curves for the largest values are still unrealistic and this method doesn’t provide a recommendation for how to split patients. A common transformation in the field is to create a binary label, and the TMB value that determines this label is viewed as the optimal cutoff (Figure 1E). However, both the Cox regression approach and the optimal cutoff approach make the assumption the risk only increases or decreases with respect to TMB. We introduce a two-cutoff approach that defines a “moderate” group and then combines the low and high groups into one “extreme” group. Depending on the data and the cutoffs selected the different cutoff approaches can give very similar answers (Figure 1F).

### Simulated Data Reveals Limitations of Current Approaches

In Figure 1 we were using actual TCGA survival data so we did not know whether the single-cutoff or two-cutoff approach was the correct way to model the data. To determine the validity of our new two-cutoff approach we can turn to simulated data. In this simulated data the TMB values are real to mimic the distribution and sample sizes we would see in actual data (the values of the UCEC TCGA samples), but the survival data is generated according to a known function mapping logged TMB values to log partial hazard. When applying a single-cutoff approach to data with a linear relationship (Figure 2A), the approach correctly identified a “TMB high” group with a significant hazard ratio for every simulation (Figure 2B).

**Figure 2.**
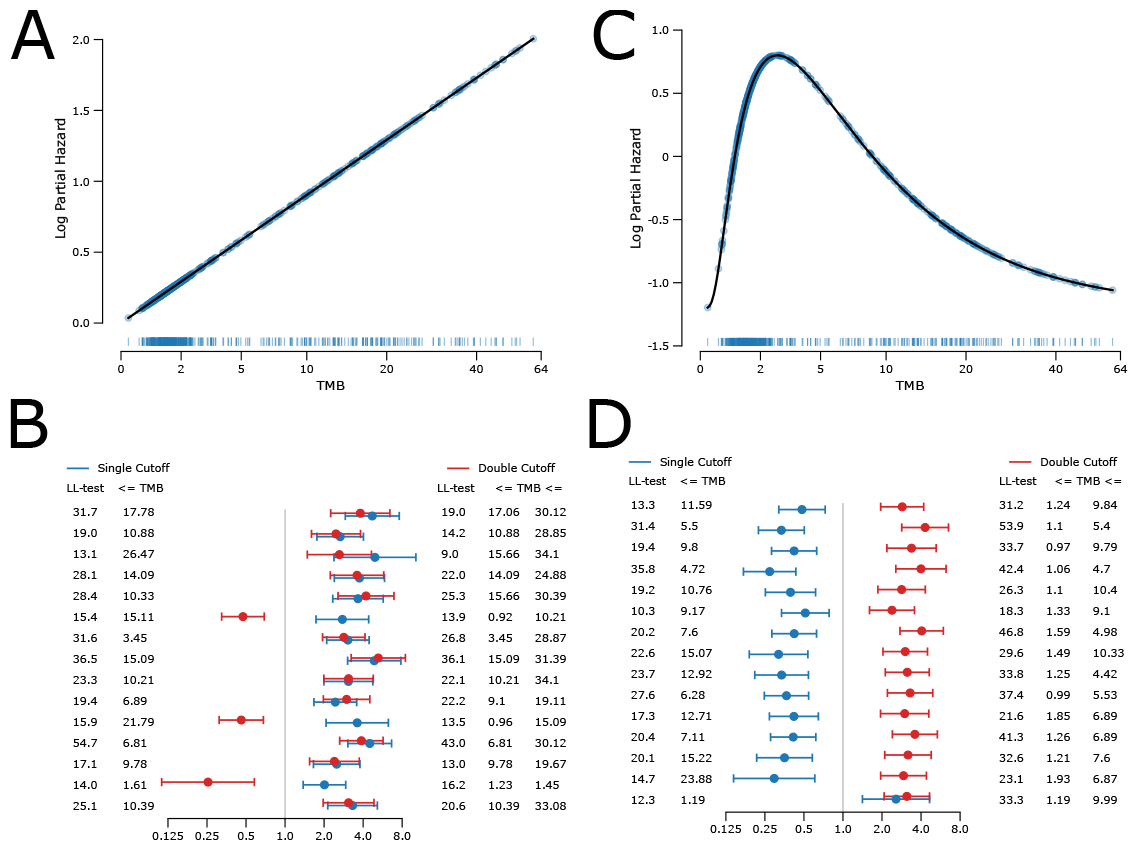
Cutoff analysis with simulated data. 15 simulated survival datasets were generated for either a monotonic relationship with TMB (A) or a non-monotonic relationship (C). B shows the hazard ratios and associated log-likelihood ratio tests and associated cutoffs of searching for an optimal cutoff or an optimal two cutoffs for a monotonic relationship, while D shows the results with non-monotonic data. In A, C the TMB distribution is shown as a rug plot with the true risks shown as a scatterplot. In B, D each row represents a different simulated data set.

Although a single-cutoff approach works well for this data, we wondered what would happen with a more complex relationship (Figure 2C). Further, we wanted to understand if our two-cutoff approach could be used to identify this type of relationship. While two cutoffs have been proposed before^27^, in those contexts three groups were generated where the “TMB mid” group was expected to have a risk in between that of “TMB low” and “TMB high”, which retains the monotonic nature of the relationship. In contrast, in our two-cutoff approach we assign the same risk to the “TMB low” and “TMB high” groups with the “TMB mid” displaying a different survival risk, which represents a non-monotonic relationship that can be described as a “Goldilocks effect” wherein we have some cases where the TMB is too “cold”, some cases where it’s too “hot”, and in between where it’s “just right”.

In the case of a monotonic relationship, we would expect grouping the highest and lowest values together would result in poor correlation, and when we applied the new two-cutoff approach to the monotonic data the statistical significance was often lower than the single-cutoff approach (Figure 2B), with the “TMB mid” group often having the higher hazard. However, in the case of a non-monotonic relationship, grouping the lowest and highest values is exactly what should be done and when these two different approaches were applied to data simulating a non-monotonic relationship of TMB with survival we found the two-cutoff approach correctly associated a middle group with a larger hazard, and always did so with greater statistical significance than the single-cutoff approach (Figure 2D).

These results give a potential framework for identifying non-monotonic relationships of an input variable to survival—if the single-cutoff approach has a better test statistic then the relationship is possibly monotonic, and if the two-cutoff approach has a better test statistic the relationship is possibly non-monotonic. However, while we generally saw this expected pattern in the simulated data it was not always the case, and this heuristic doesn’t reveal the true underlying relationship, such as a linear relationship versus a step function or others. Instead of comparing different transformation strategies (one cutoff versus two), we can simply allow a fully connected neural network (FCN) to learn the relationship between predictor variable and outcome measures directly from the available data.

We looked at the first two simulated survival datasets for both the linear and non-monotonic relationships, and compared the results of a fully connected network (FCN) to a Cox regression implemented with lifelines. To compare the model fits we looked at the C-index and log-likelihood (a larger value is better in both cases) for data in the test folds of 10 stratified K-folds, and also compared these metrics to what would be obtained with the ground truth relationship (Table 1). With regards to the linear data both the Cox model and the FCN had metrics nearly indistinguishable from the ground truth, but for the non-monotonic data the FCN had noticeably better metrics than the Cox model.

**Table 1.** Simulated data metrics. Log-likelihood and C-indexes for the test folds of two representative monotonic and non-monotonic simulated datasets from Figure 2 for either a Cox model or a neural network.

| Simulated Dataset | Model | Linear Data |  | Non-monotonic Data |  |
| --- | --- | --- | --- | --- | --- |
|  |  | LL score | C-index | LL score | C-index |
| 1 (Row 1) | Ground Truth | -.895 | .601 | -.905 | .651 |
|  | Cox-PH | -.895 | .601 | -.929 | .528 |
|  | FCN | -.896 | .601 | -.909 | .646 |
| 2 (Row 2) | Ground Truth | -.921 | .620 | -.872 | .665 |
|  | Cox-PH | -.921 | .620 | -.892 | .580 |
|  | FCN | -.922 | .620 | -.884 | .645 |

We also visualized the model fits of the FCNs, Cox regressions, and the true underlying relationships (Figure 3). Despite having a large number of parameters our FCN produced a very similar fit to a Cox regression for the linear data, with both closely following the true relationship (Figure 3A, B). Visually, the extra parameters of the FCN allowed it to correctly follow the shape of the non-monotonic data (Figure 3C, D), while the Cox model was a poor fit to the data, which is consistent with the model metrics. For additional data distributions see Supplemental Figure 1. For the variability in model fits across training folds see Supplemental Figure 2.

**Figure 3.**
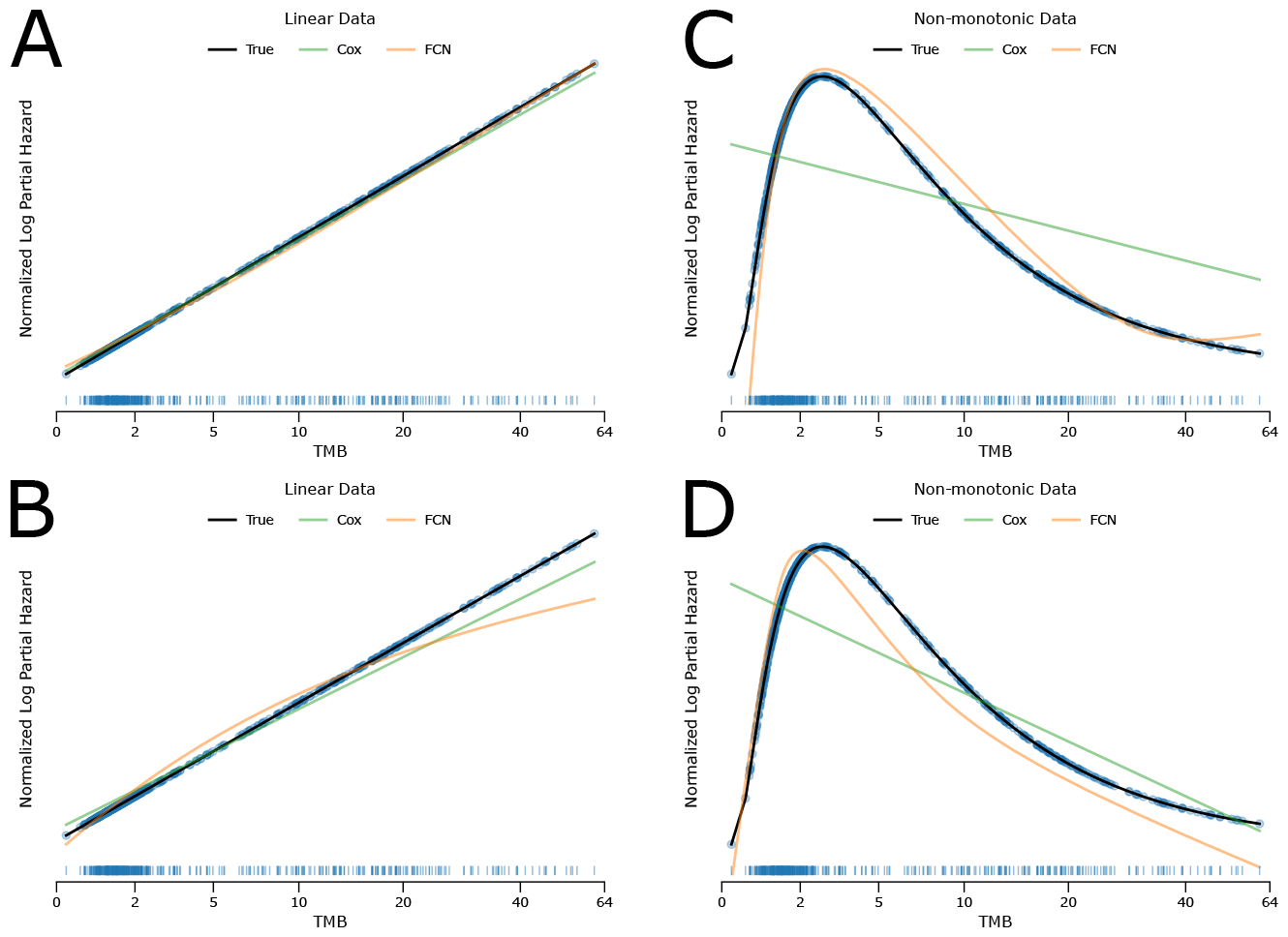
Simulated data model fits. Mean-normalized model fits from a Cox model and a neural net were averaged over 10 K-folds for the first two simulated survival datasets for a linear and non-monotonic relationship. A, B monotonic data. C, D non-monotonic data. TMB distributions shown as rug plots with the true risks shown as a scatterplot.

### Applying Neural Networks to the TCGA

The simulated data gave us confidence that if the relationship strongly deviates from a monotonic relationship, then an FCN will be able to better model this and this would be reflected in the C-index and model log-likelihood metrics. Our next question was whether this tool would allow us to find evidence for such “Goldilocks effects” in real-world datasets.

Using the mutation call and survival data for solid tumors in the TCGA dataset, we compared the model metrics of a Cox model and our FCN (Table 2).

**Table 2.** TCGA data metrics. Log-likelihood scores and C-indexes for the test folds of different cancers in the TCGA for a Cox model and a neural net.

| Cancer | Cox-PH |  | FCN |  |
| --- | --- | --- | --- | --- |
|  | LL score | C-index | LL score | C-index |
| BLCA | -1.337 | .573 | -1.338 | .570 |
| CESC | -.593 | .512 | -.600 | .490 |
| ESCA | -.842 | .556 | -.842 | .556 |
| GBM | -1.967 | .491 | -1.967 | .478 |
| HNSC | -1.381 | .532 | -1.379 | .540 |
| KIRC | -.643 | .616 | -.643 | .616 |
| KIRP | -.372 | .573 | -.372 | .573 |
| LGG | -.630 | .712 | -.625 | .712 |
| LIHC | -.953 | .572 | -.955 | .572 |
| LUAD/LUSC | -1.488 | .514 | -1.488 | .514 |
| OV | -1.415 | .569 | -1.416 | .569 |
| PAAD | -1.177 | .551 | -1.193 | .553 |
| COAD/READ | -.670 | .580 | -.670 | .580 |
| SKCM | -1.346 | .578 | -1.322 | .617 |
| STAD | -1.203 | .559 | -1.204 | .559 |
| UCEC | -.515 | .551 | -.521 | .562 |

In most cancer types there was limited difference between a Cox model and an FCN, suggesting the relationship likely is monotonic (or in some cases simply no relationship). However, in SKCM the FCN displayed a noticeably better log likelihood and C-index. SKCM is a cancer type where immunotherapy is recommended and where TMB has been suggested as predictive biomarker^28,29^. Looking at some of the model fits we see tumor types where the FCN predicted a perfectly linear relationship and followed the Cox predictions, cases where the FCN has slight deviations from the Cox predictions, and then SKCM where the FCN predicts a strong concave up relationship (Figure 4).

**Figure 4.**
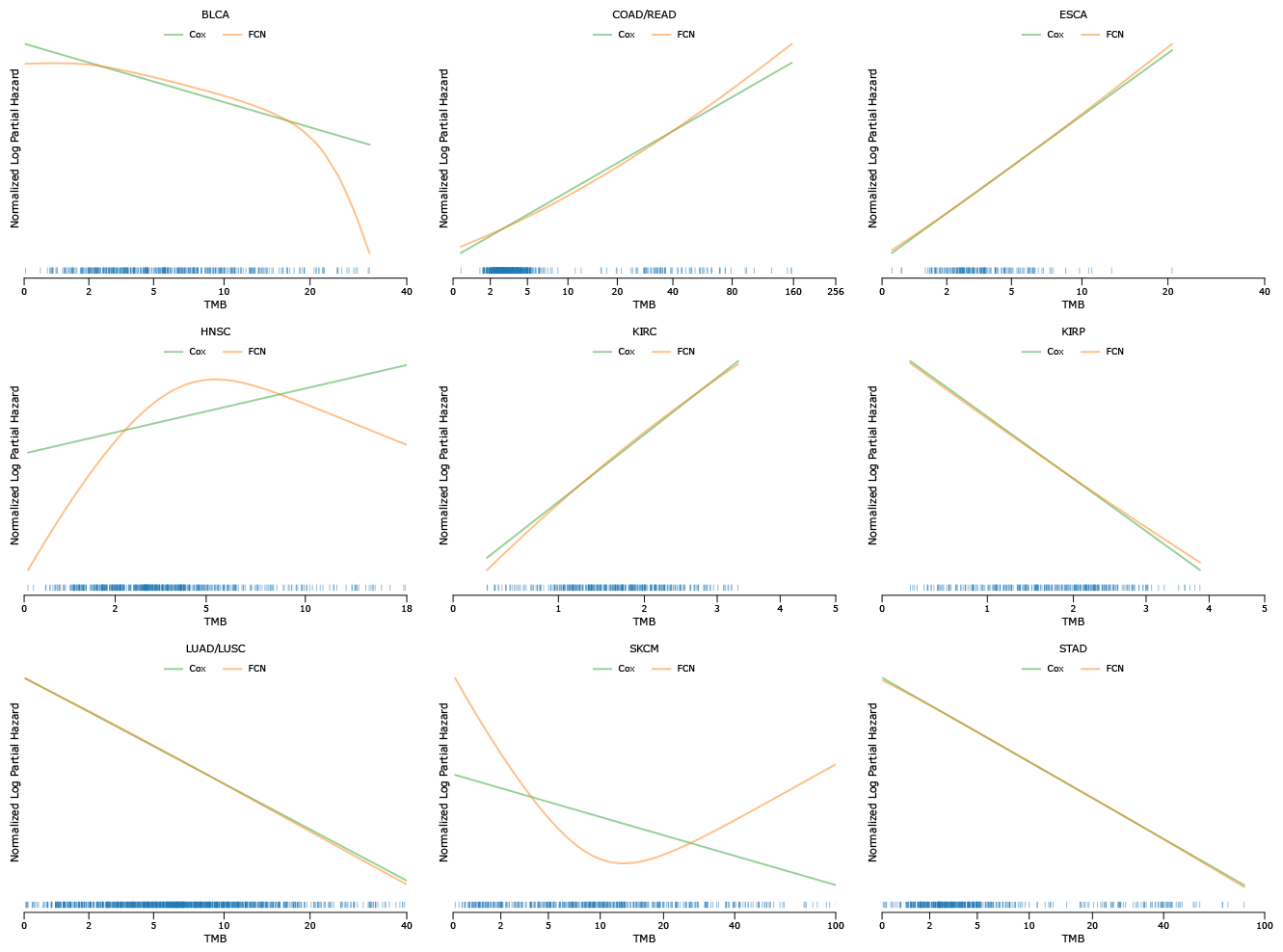
TCGA model fits. Nine selected cancer types are shown. Mean-normalized Cox and neural net model fits were averaged over 10 K-folds. TMB distributions shown as rug plots.

### Applying Neural Networks to Immunotherapy Datasets

The TCGA dataset contains patients receiving the standard of care at the time, and includes patients at different stages of disease. In this heterogeneous context, exploring the relationship of TMB with survival is more closely associated with some form of prognostication rather than predicting response to a given therapeutic modality. In contrast, when investigating a cohort in which a specific treatment has been administered we generally are considering a biomarker that is predictive of response. It is in this context that TMB has often been explored as a biomarker, and many approaches effectively attempt to stratify the cohort into “TMB low” and “TMB high” and compare clinical outcome measures (response, disease-free survival, etc). Given our findings of a non-monotonic relationship of survival for SKCM in the TCGA dataset we wondered if we would identify non-monotonic relationships in other datasets, either in a prognostic or predictive context.

The AACR Project GENIE Biopharma Collaborative (BPC) recently released clinical data for lung and colon cancer patients. Using the corresponding panel mutational data for these patients we applied our model to a lung cohort treated without immunotherapy (BPC NSCLC NonIO), a lung cohort treated with immunotherapy (BPC NSCLC IO), and a colon cohort treated without immunotherapy (BPC colorectal NonIO). Similar to the TCGA data our neural network did not detect non-monotonic relationships in these cancer types (Table 3, Supplemental Figure 3).

**Table 3.**
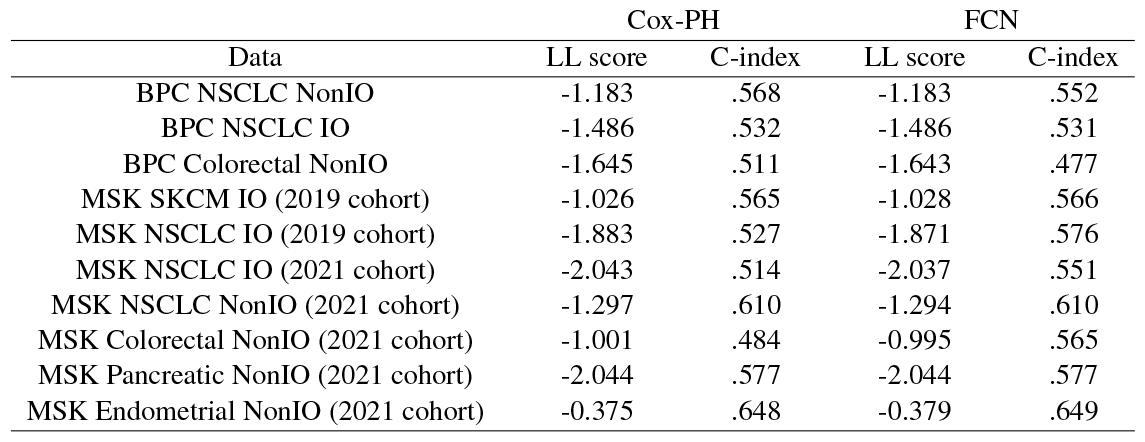
BPC and MSK data metrics. Log-likelihood scores and C-indexes for the test folds of different cancers in the BPC and MSK data for a Cox model and a neural net.

Memorial Sloan Kettering (MSK) has released several large datasets of patients assayed with their gene panel and the corresponding clinical information. Without some form of common identifier across these datasets it’s difficult to know how much patient overlap exists between the data releases, but we applied our FCN to a 2019 dataset which only included patients treated with immunotherapy^15^, and a 2021 dataset that had both IO naive and IO treated patients^18^. While we had observed TMB to have a non-monotonic relationship with survival in the TCGA dataset for SKCM, in the IO treated MSK melanoma cohort we did not observe a difference between a Cox regression and our FCN (Table 3). This is consistent with a monotonic relationship of TMB and clinical outcome in the context of melanoma treated with IO, and supports a TMB-high vs TMB-low stratification in this context. The FCN model showed better performance over Cox modeling in the non-IO treated colorectal cancer cohort (MSK Colorectal NonIO), in which a concave down model was apparent (Supplemental Figure 4). Some improvement in performance was seen in the MSK non-small cell lung cancer cohorts treated with IO, and the observed relationship was also concave down (Supplemental Figure 4).

### Example of Model Application

The benefit of directly modeling the relationship with survival is that any deviations from a linear fit will be accounted for in the model predictions. This allows a researcher wanting to identify a cutoff for splitting patients to simply use a cutoff based on model risks. Then, working backwards from the model risks the corresponding input variable cutoff(s) can be found. If the relationship of the input variable to survival is non-monotonic then a single model risk will result in two input variable values, while a monotonic relationship will result in a single input variable cutoff.

We can demonstrate what this might look like by splitting each cancer in the TCGA dataset by either the median TMB value or median FCN model risk value. When splitting by median TMB the higher TMB group may or may not have a higher hazard, while when splitting by median model risk the higher risk group should have a higher hazard. Ignoring this difference in sign, in most cases there is almost no difference in the test statistic since the FCN predicted a monotonic relationship for most cancers (Figure 5A, B). However, for SKCM we see that a single cutoff of TMB is inappropriate. In fact, with a median cutoff the relationship of TMB to survival is barely significant while it is highly significant with a FCN model output-based risk cutoff (Figure 5C, D).

**Figure 5.**
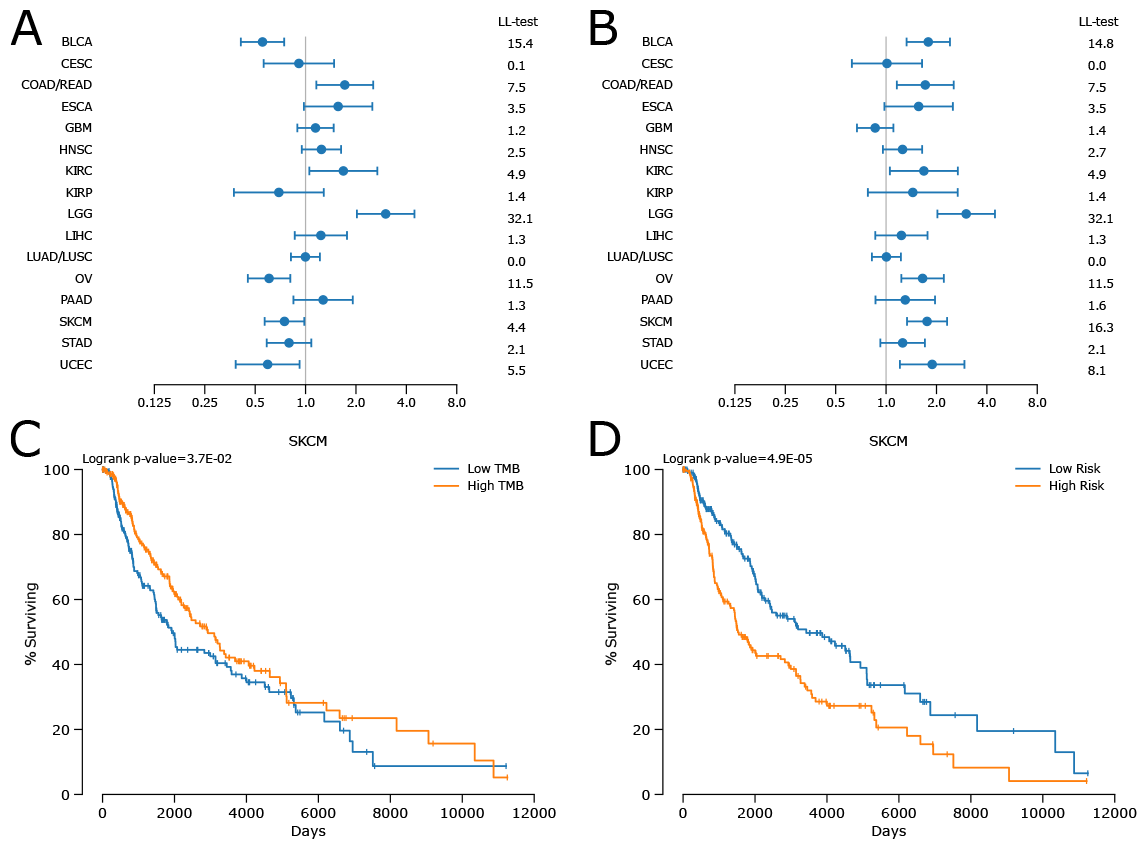
Neural network can create a better binary label. Patients were split by either median TMB (A, C), or median risk from a neural net (B, D). A, B show the hazard ratios and log-likelihood tests of each cancer type for these splits while C, D show the Kaplan curves for SKCM with these splits along with the logrank p-value.

## Discussion

Identifying consistent cutoffs for biomarkers has always been challenging^30^ since only relationships that resemble a step function result in stable optimal cutoff values^31^, a result recapitulated in our Figure 2 and Supplemental Figure 1. It is possible to identify a step function with a neural net (Supplemental Table 1, Supplemental Figure 1), but here we used neural nets to investigate yet another consideration when identifying potential biomarkers and cutoffs: the assumption of a monotonic relationship may not hold. If this scenario occurs it has several important implications. First, when no relationship is found with a Cox regression a very strong relationship could still be present. Second, the predictions for values at the extremes, i.e. patients who would be predicted to have the best or worst survival according to a Cox model, will contain the largest errors (see the fits for SKCM in Figure 4). Third, if a non-monotonic relationship is present then a single cutoff is inappropriate and two-cutoffs of the input variable will be needed to split patients into the appropriate two groups.

The possibility of a non-monotonic relationship is not just theoretical. While approaches to correlate TMB to survival have generally assumed a monotonic relationship, using a more flexible approach we have identified multiple datasets where the relationship between TMB and survival appears more complex. This study has a few important limitations, the first of which is that we did not undertake a more complex multivariate modeling of clinical outcome using features such as age, stage, grade, etc. This was done on purpose as the intent of this work is to highlight how the relationship of TMB to clinical outcome can be modeled to account for potential non-monotonic relationships and not to comprehensively approach clinical outcome modeling across the relevant predictors. The second important limitation is related in that we did not design these experiments such that we would be “validating” a specific TMB *↔* clinical outcome correlation across tumor type and clinical setting. Again, this was intended given the modeling focus of the work. We believe what we have developed and presented herein has important implications for studies that investigate TMB as a predictive and/or prognostic biomarker. Further, we show that fairly simple neural networks like we have presented here can help avoid the pitfalls of ordinary Cox-PH based regression with respect to the potential of non-monotonic relationships.

## ACKNOWLEDGMENTS

The results here are in whole or part based upon data generated by the TCGA Research Network. The authors would like to acknowledge the American Association for Cancer Research and its financial and material support in the development of the AACR Project GENIE registry, as well as members of the consortium for their commitment to data sharing. Interpretations are the responsibility of study authors.

