## Supplemental Information for "Characterization of non-monotonic relationships between tumor mutational burden and clinical outcomes"

| Model | LL score | C-index |
|---|---|---|
| Ground Truth | -.944 | .629 |
| Cox-PH | -.963 | .619 |
| FCN | -.958 | .618 |
| Sigmoid | -.951 | .619 |

**Supplemental Table 1. Simulated step function metrics.** Log-likelihood and C-indexes for the test folds of a simulated step function dataset for either a Cox model, a FCN neural network, and a neural network comprised of a single neuron with sigmoid activation.

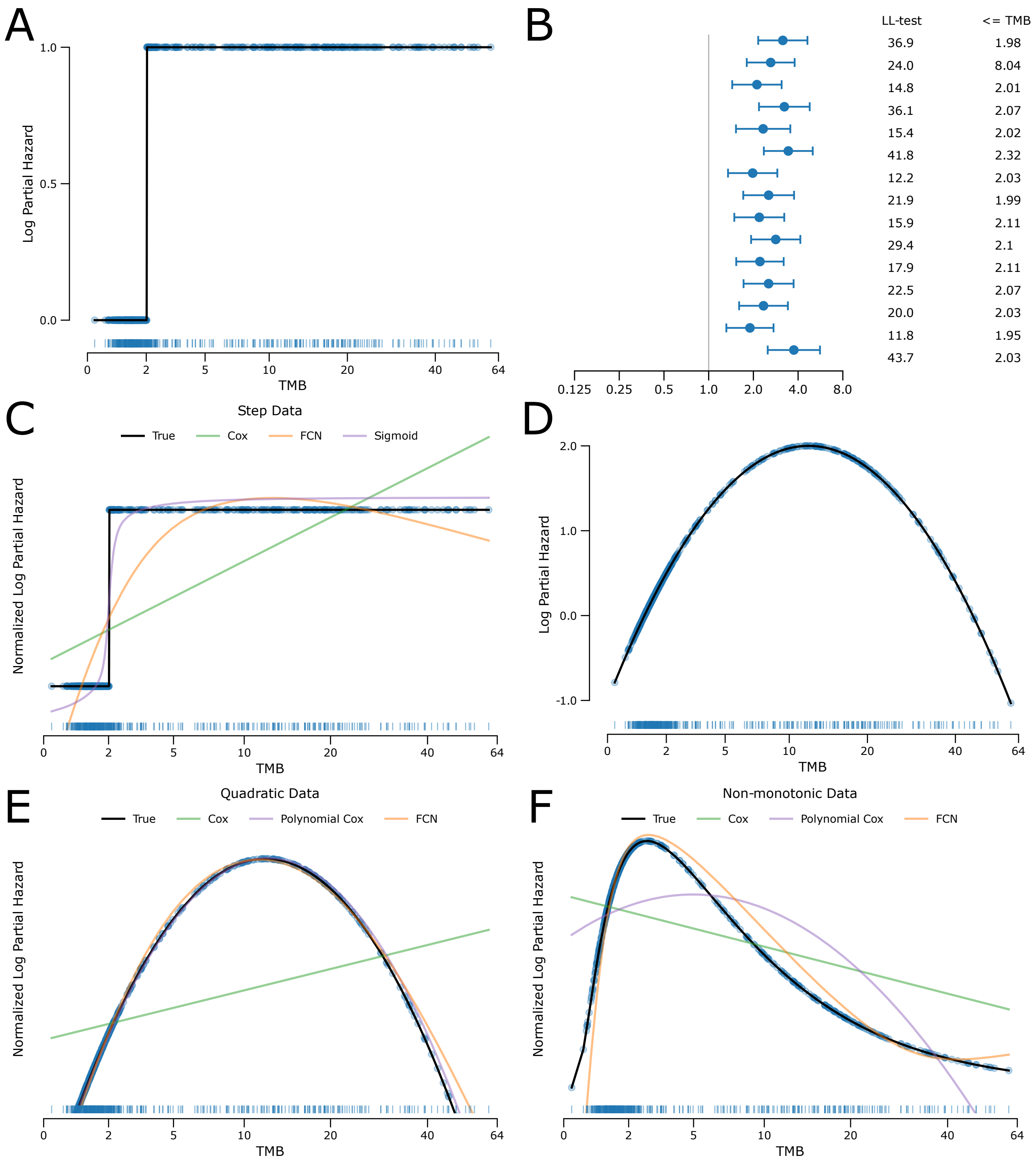


**Supplemental Figure 1. Simulated step data and examples with a polynomial transformation.** 15 simulated survival datasets were generated for a step relationship with TMB (A). B shows the hazard ratios and associated log-likelihood ratio tests and associated cutoffs of searching for an optimal cutoff, while C shows the fits of a Cox model, FCN neural network, and a neural network comprised of a single neuron with sigmoid activation. D shows an example of a quadratic relationship with TMB (E). Generating a simulated dataset from the risk relationship in D we explored fitting a Cox model with a two degree polynomial in addition to a neural net. In F we show the fit of a two degree polynomial with non-monotonic data.

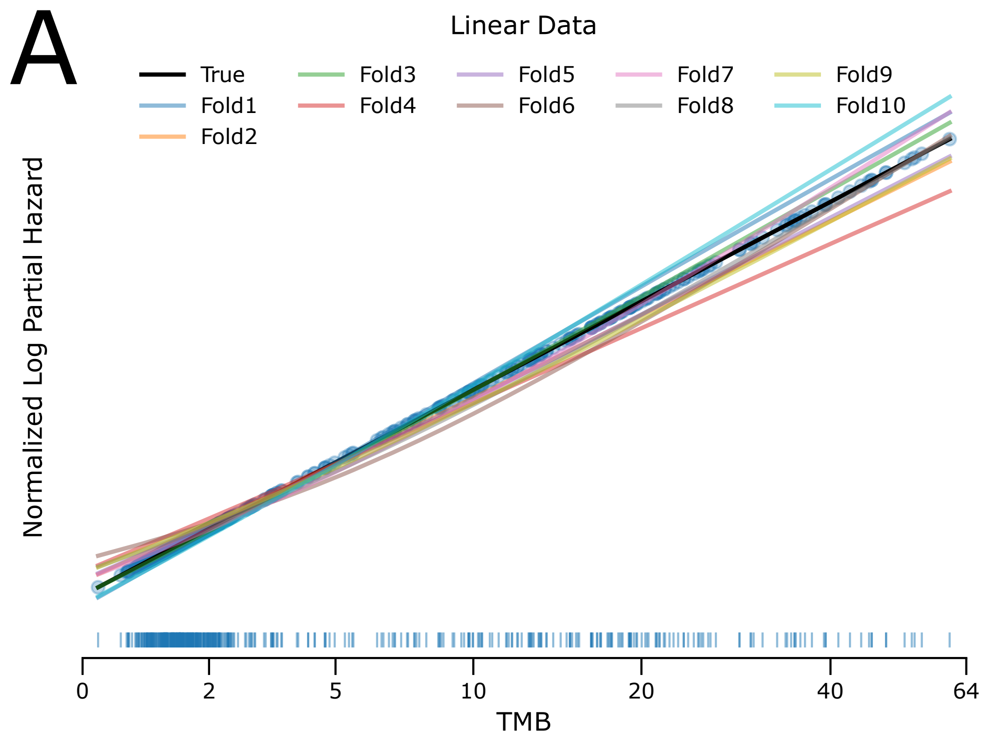


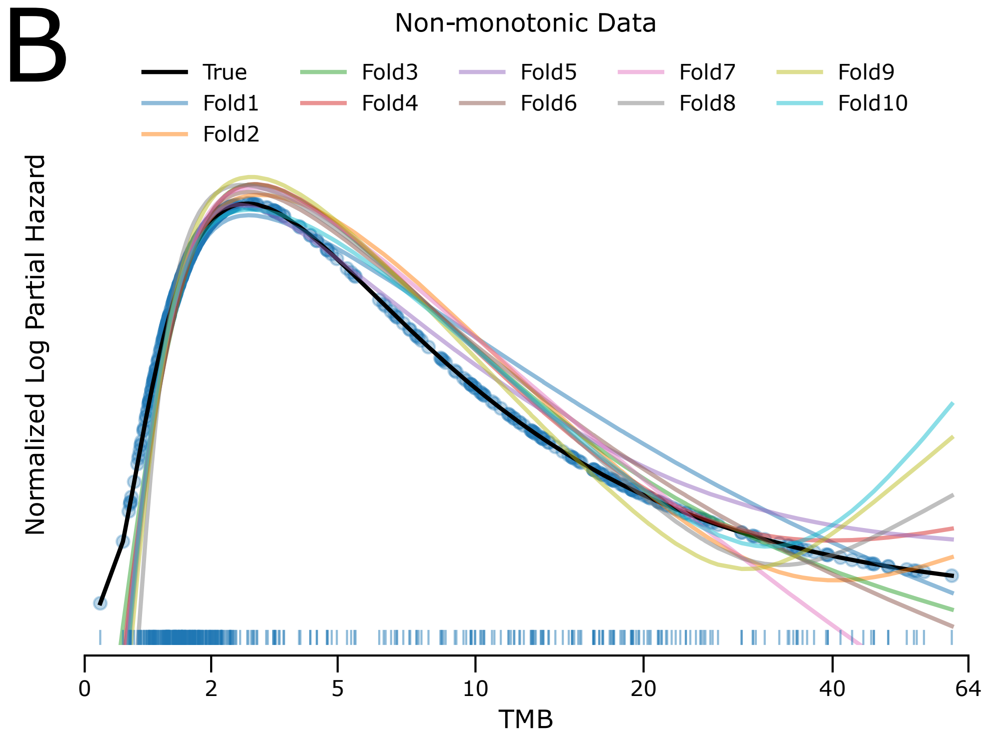


**Supplemental Figure 2. Model variability across folds.** A, mean-normalized model fits for each training fold for an FCN model with simulated linear data. B, mean-normalized model fits for each training fold for an FCN model with simulated non-monotonic data.

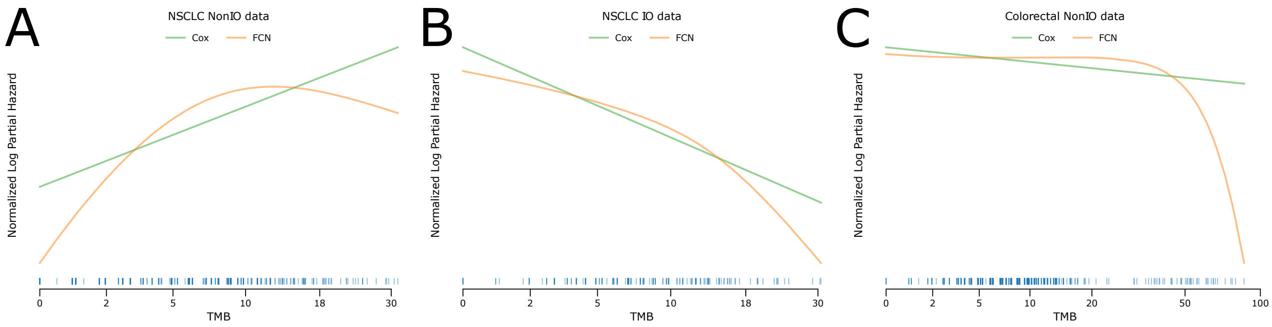


**Supplemental Figure 3. BPC fits.** Cox and neural net model fits were mean normalized and averaged over 10 K-folds. TMB distributions shown as rug plots.

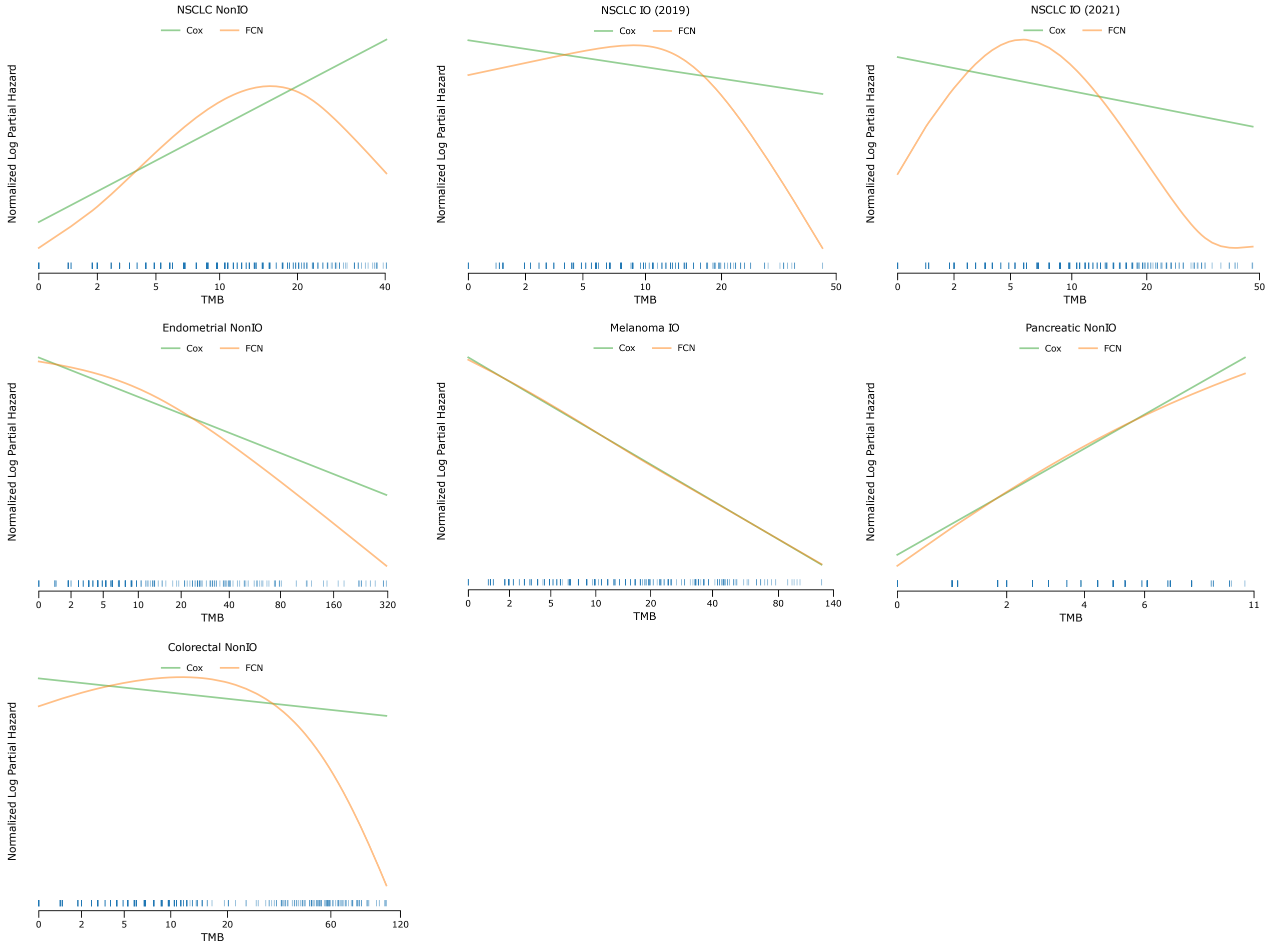


**Supplemental Figure 4. MSK fits.** Cox and neural net model fits were mean normalized and averaged over 10 K-folds. TMB distributions shown as rug plots.
